# A New Era in Missense Variant Analysis: Statistical Insights and the Introduction of VAMPP-Score for Pathogenicity Assessment

**DOI:** 10.1101/2024.07.11.602867

**Authors:** Eylul Aydin, Berk Ergun, Ozlem Akgun-Dogan, Yasemin Alanay, Ozden Hatirnaz Ng, Ozkan Ozdemir

## Abstract

The clinical interpretation of missense variants is critically important in diagnostics due to their potential to cause mild-to-severe effects on phenotype by altering protein structure. Evaluating these variants is essential because they can significantly impact disease outcomes and patient management. Many computational predictors, known as in silico pathogenicity predictors (ISPPs), have been developed to support the assessment of variant pathogenicity. Despite the abundance of these ISPPs, their predictions often lack accuracy and consistency, primarily due to limited data availability and the presence of erroneous data. This inconsistency can lead to false positive or negative results in pathogenicity evaluation, highlighting the need for standardization. The necessity for reliable evaluation methods has driven the development of numerous ISPPs, each attempting to address different aspects of variant interpretation. However, the sheer number of ISPPs and their varied performances make it challenging to achieve consensus in predictions. Therefore, a comprehensive statistical approach to evaluate and integrate these predictors is essential to improve accuracy. Here, we present a comprehensive statistical analysis comparing 52 available ISPPs, which aims to enhance the precision of variant classification. Our work introduces the Variant Analysis with Multiple Pathogenicity Predictors-score (VAMPP-score), a novel statistical framework designed for the assessment of missense variants. The VAMPP-score leverages the best gene-ISPP matches based on ISPP accuracies, providing a combinatorial weighted score that improves missense variant interpretation. We chose to develop a statistical framework rather than creating a new ISPP to capitalize on the strengths of existing predictors and to address their limitations through an integrative approach. This approach not only improves the evaluation of missense variants but also offers a flexible statistical framework designed to identify and utilize the best-performing ISPPs. By enhancing the accuracy of genetic diagnostics, particularly in the reanalysis of rare and undiagnosed cases, our framework aims to improve patient outcomes and advance the field of genetic research.

Our study employed a comprehensive workflow (Figure 1) to enhance the accuracy of genomic variant interpretation with in-silico pathogenicity predictor (ISPP) evaluation. This workflow led to three pivotal results:

● ISPPs were categorized on their prediction approaches. This classification not only streamlined the analytical process but also enhanced the interpretability of predictor outputs.
● Leveraging this categorization, we conducted a robust statistical analysis to evaluate the prediction accuracy and performance of each ISPP. Our findings revealed a significant correlation between the prediction approaches of the ISPPs and their predictive successes, confirming the utility of our categorization approach.
● These insights enabled us to develop a novel scoring system—the VAMPP-score—which integrates ISPPs according to their performances.

---

**Figure 1.**
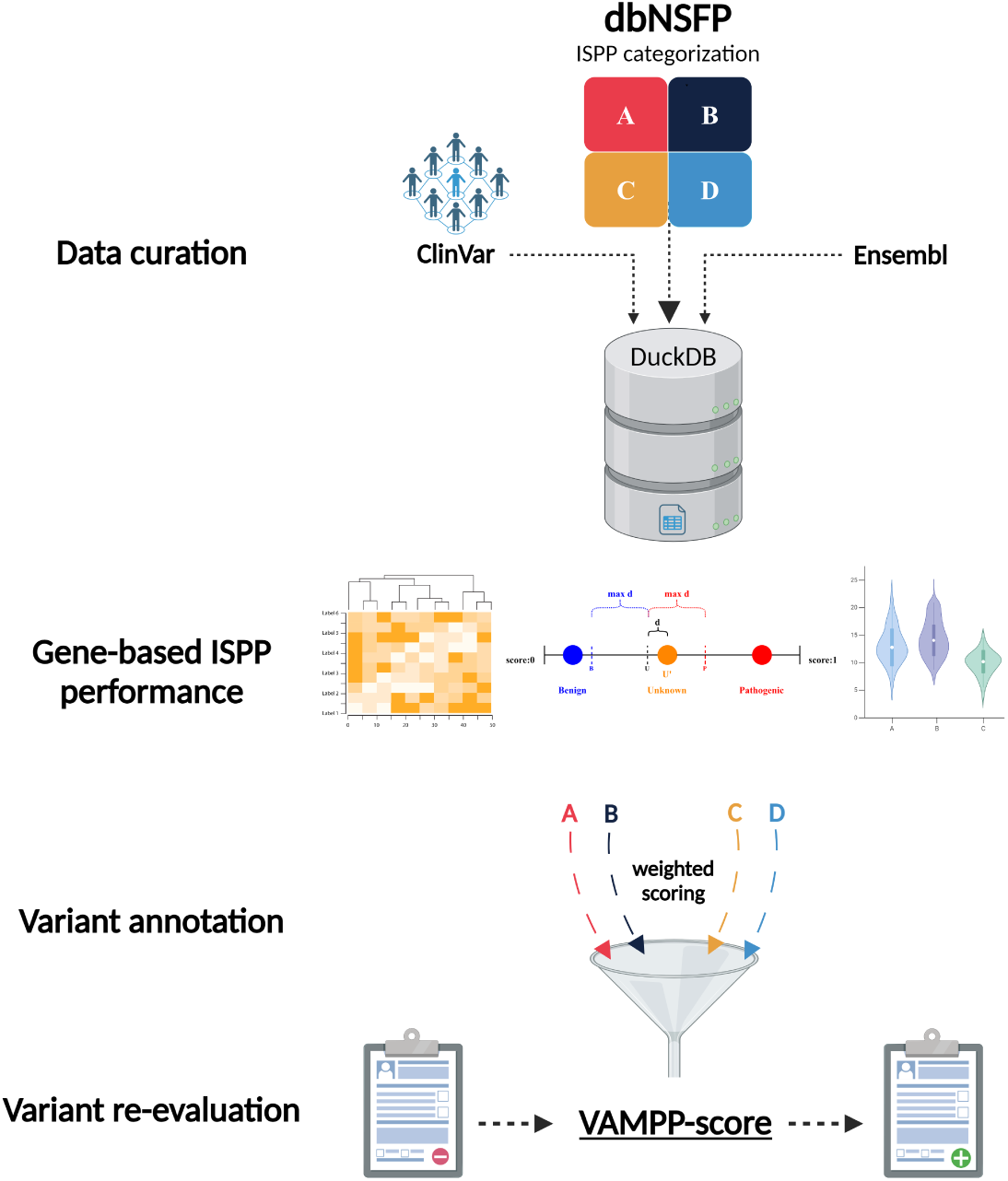
Methodological summary. A database was built using dbNSFP (1, 2) (v4.7A) data, which includes rank scores from over fifty computational predictors for all possible single nucleotide variations in the human genome. This database was enhanced with canonical transcript information from Ensembl. The analysis focused on missense variants in ClinVar (3), categorized as Pathogenic (P), Benign (B), and Unknown (U). A novel statistical framework was used to assess the gene-based performance of each ISPP in distinguishing these variant groups through multiple and pairwise comparisons and proximity estimations. ISPPs were categorized (Category A-D) and assigned weights based on their performance for each gene, known as gene coefficients. These were integrated into the final assessment, the VAMPP-score.

## 1 Introduction

Advances in sequencing technologies, particularly next-generation sequencing (NGS), have significantly transformed the field of genetic diagnosis, enabling comprehensive, rapid, and cost-effective genetic analyses that enhance our understanding of genetic disorders. The significant reduction in sequencing costs has enabled easier access to genetic testing, broadening its availability across a wider range of patients and clinical settings (4).

Mendelian disorders, identifiable by their unique inheritance patterns, constitute a significant portion of the genetic conditions detected through NGS-related technologies (5, 6). These disorders, often caused by disruptions in single genes, have provided critical insights into the genetic basis of various diseases. With NGS, researchers have identified novel variants in known disease-causing genes and discovered new genes associated with these conditions despite the complexities posed by genetic interactions and the phenotypic heterogeneity observed even within the same disorder (6, 7).

One of the most encountered and challenging variations is the missense variants, which involve amino acid changes through single nucleotide substitutions. These variants are notoriously difficult to interpret due to their subtle yet significant impacts on protein function, disrupting protein structure and function and consequently influencing biological phenotypes (8). Real-world clinical settings rarely allow for conclusive functional studies to determine their pathogenicity, making reliance on computational pathogenicity predictors necessary (9).

However, these in-silico tools often yield conflicting and inaccurate predictions, leading to false positives and negatives that can significantly affect patient diagnosis and management (10). Recognizing these limitations, the American College of Medical Genetics and Genomics (ACMG) and the Association for Molecular Pathology (AMP) have established comprehensive guidelines, including the PP3 criterion, which calls for the agreement of multiple computational evidence to support the pathogenicity of a variant (11). These guidelines have substantially improved the consistency and accuracy of genetic diagnoses but still face challenges, particularly with rare genetic disorders where less information is available (5, 12).

Another significant challenge in genetic diagnostics is dealing with Variants of Uncertain Significance (VUS), whose health impacts are poorly understood. This issue is exacerbated in the context of rare diseases, where many genes and genetic variants remain under-studied and poorly documented, complicating clinical decision-making and raising ethical questions about how uncertainties should be communicated to patients (11, 13–15).

To address these challenges, we analyzed the prediction accuracy of the most frequently used ISPPs using statistical approaches to strengthen variant interpretation in diagnostics. This approach led us to develop the Variant Analysis with Multivariate Pathogenicity Prediction-Score (VAMPP-score), a novel in-silico pathogenicity prediction score for missense variants. This score employs a statistical framework that utilizes multiple and pairwise comparison tests, proximity estimations, and mean difference calculations among variant groups. It also leverages gene-specific ISPP accuracies to provide a combinatorial weighted score. The VAMPP-score is designed to be a valuable tool for clinicians in the diagnosis of genetic disorders, particularly for the reanalysis of rare and previously undiagnosed cases, promising to increase the success rate of diagnoses and enhance the accuracy and effectiveness of genetic diagnostics, ultimately improving patient outcomes.

## 2 Results

### 2.1 In-silico pathogenicity predictors (ISPPs) considered

In this study, we enrolled a total of 52 ISPPs and categorized them according to their prediction approaches into four groups (Figure 2) (Supplementary Table 1). These groups were shaped after a comprehensive literature review and creating a sensible ground, with an effort not to generalize the aspects while trying to categorize under main features.

**Figure 2.**
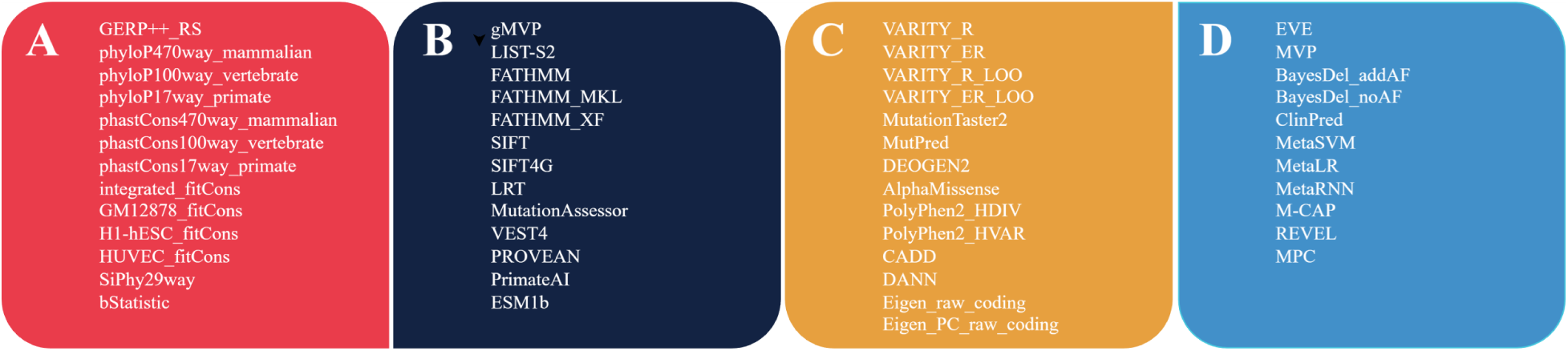
ISPP categorization. Based on their prediction approaches, the ISPPs with available ranked scores in dbNSFP4.7A (1, 2) were categorized into four groups (A, B, C, and D).

*Category A* ISPPs utilize evolutionary conservation at the nucleotide sequence level, using multiple sequence alignment (MSA) to identify functional regions affected by genome variations. Tools like phyloP (16), phastCons (17), and GERP++ (18) fall into this category. *Category B* ISPPs focus on amino acid sequence conservation, primarily for missense variants such as. SIFT (19), which predicts damaging changes at conserved amino acid positions. Other tools include FATHMM (20), PROVEAN (21), and VEST4 (22). *Category C* ISPPs, known as meta-predictors, integrate multiple parameters such as evolutionary conservation, biological function, protein structure, and population frequency data. Tools like CADD (23), which uses a linear kernel support vector machine (SVM), DANN (24), which enhances CADD’s approach with a deep neural network, and AlphaMissense (25), noted for its proteome-wide missense variation analysis using Alphafold2 (26), exemplify this category. *Category D* ISPPs combine various parameters and scoring algorithms to enhance prediction accuracy. Tools like ClinPred (27) falls into this category, as it integrates machine learning models and 16 ISPP scores to assess the clinical relevance of variants, utilizing allele frequency data from gnomAD (28).

### 2.2 Overview

A total of 1,162,673 variants from canonical transcripts (3, 29) (Supplementary Figure 1, Supplementary Table 2) of 17,712 unique genes were analyzed. Analysis revealed that 3,955 genes had at least one ISPP with a significant result (p<0.05) in multiple comparisons (30). Furthermore, a “Mendeliome variant list” to analyze the variants in disease-causing genes based on the human phenotype ontology (HPO) (31–33) (HP:0034345). The analysis of 841,674 variants (Supplementary Table 2) of 4,703 genes resulted in 3,193 genes with significant ISPPs.

Consistent performance of ISPPs was shown across both exome (Supplementary Figure 2) and Mendeliome (Figure 3), with a shared top eight best-performing ISPP list, confirming their robustness in variant interpretation for a higher number of genes. MetaRNN (34), ClinPred (27), and BayesDel_addAF (35) emerged as the top performers (Supplementary Table 3).

**Figure 3:**
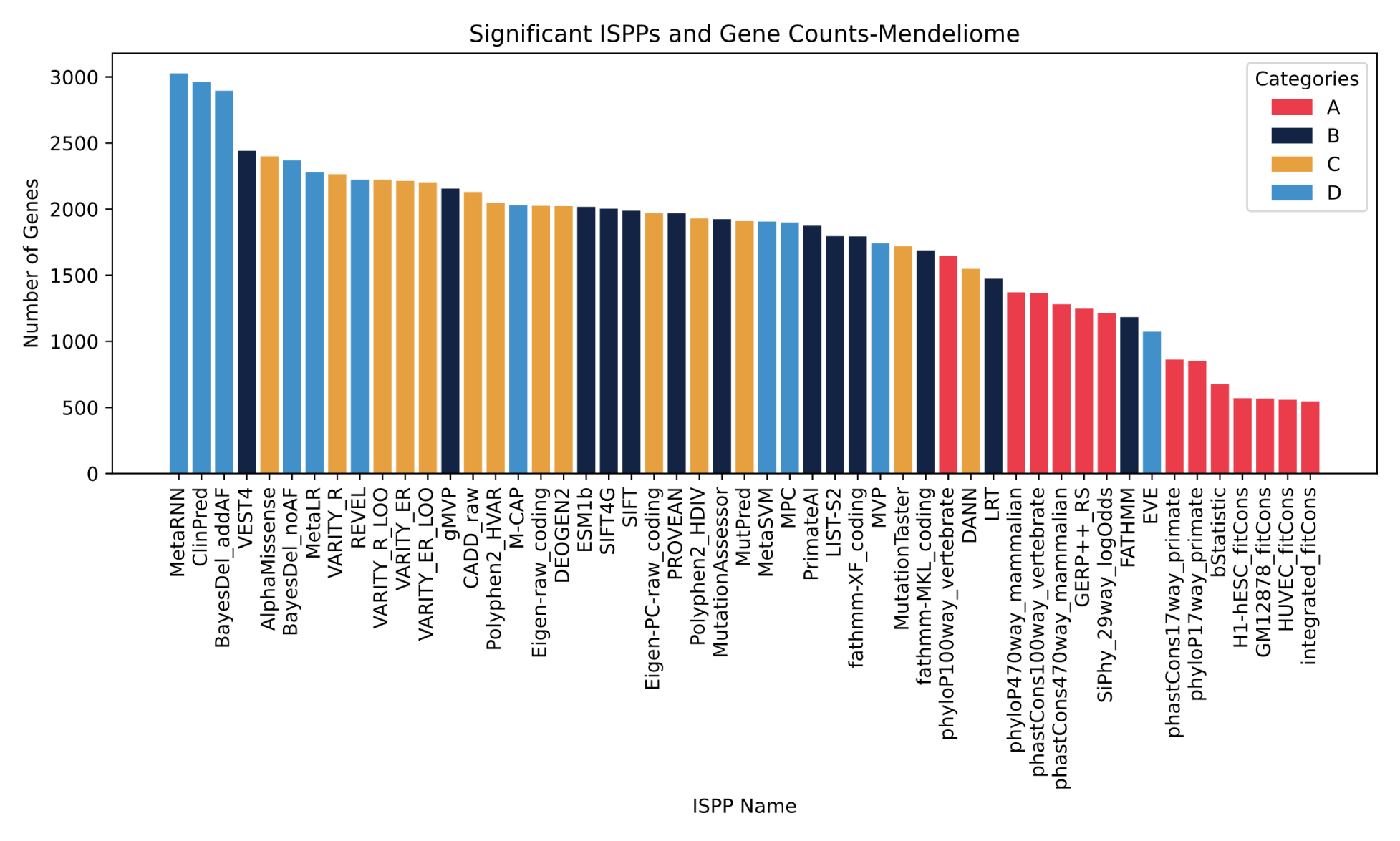
The bar plot shows the ISPPs analyzed on the x-axis and gene counts on the y-axis. Gene counts represent the number of genes each ISPP found to be significant (p<0.05) in multiple comparisons for the Mendeliome gene list. The bars in the plot are colored based on ISPP categorization, shown in the color legend.

It was noted that the top eight ISPPs (Figure 3), predominantly belong to categories C and D, except VEST4 (22) from category B. VEST4, trained on exome data, identifies rare missense variants with high disease-causing potential. While its main focus is missense variant classification, VEST4’s gene-level analysis aligns with the ensemble ISPPs of category D, ensuring predictive consistency. The leading ISPPs—MetaRNN (34), ClinPred (27), and BayesDel_addAF (35)—integrate multiple scores from across other categories, enhancing their pathogenicity predictions and affirming the categorization made and the efficacy of ensemble predictors.

### 2.3 Multidimensional Analysis

Expectations were for ISPPs of category A to exhibit distinct clustering due to their narrow approach and PCA results revealed these ISPPs as the most dispersed within their category, despite showing distinct clustering. Interestingly, ISPPs of the fitCons (37) algorithm are clustered together but separately from others in category A. This suggests that employing different cell lines for variant effect prediction did not improve performance, as evidenced by fitCons’ lower predictive accuracy (Figure 3), highlighted in the clustering results (Figure 4A).

**Figure 4:**
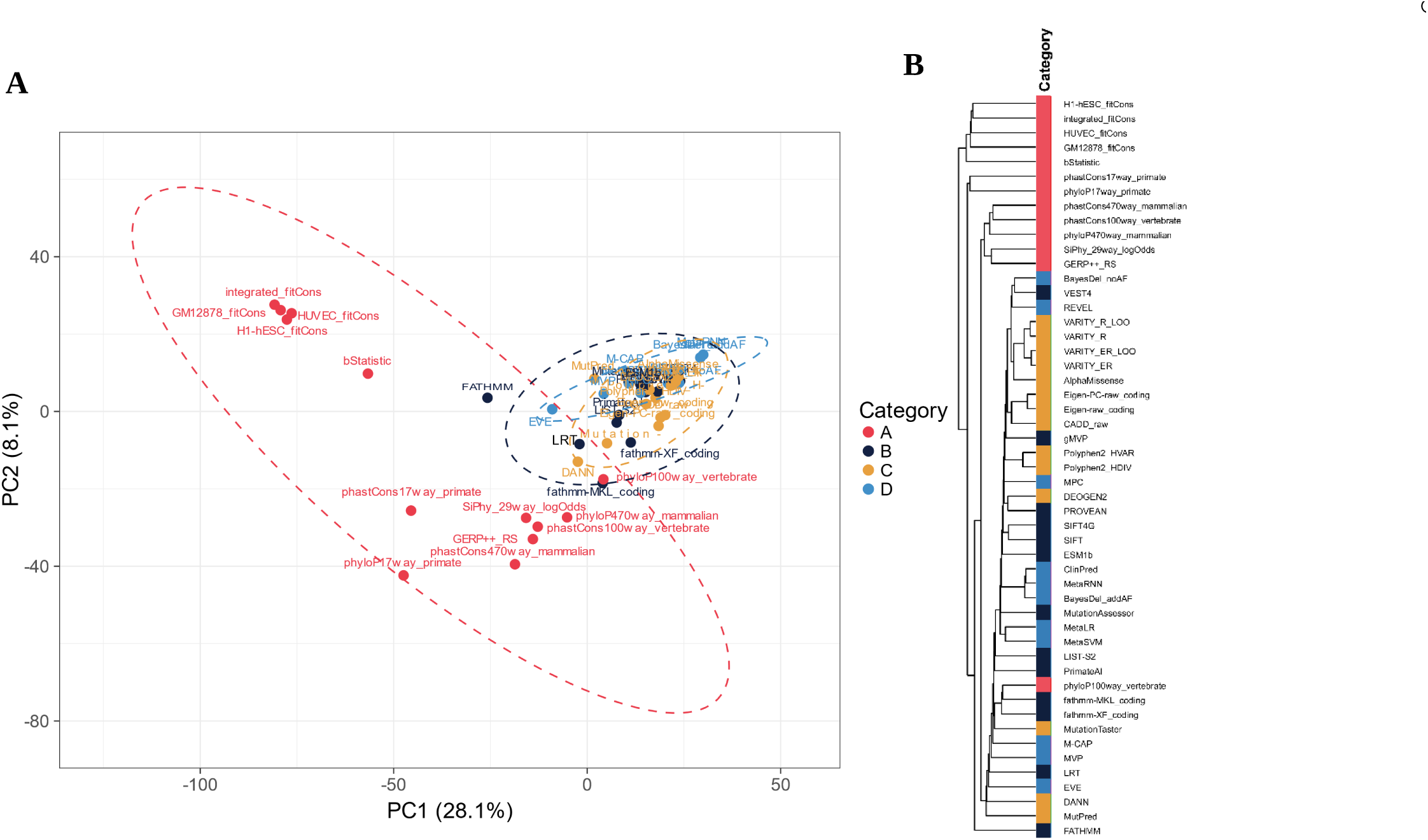
**A**. Principal component analysis (PCA) for the Mendeliome demonstrates clustering according to p-values of ISPPs from multiple comparison (30), with each category represented by differently colored dotted ellipses as indicated in the legend. The analysis was based on the first two principal components (PC1 and PC2), with additional details provided in a scree plot in the supplementary materials (Supplementary Figure 3). **B.** Hierarchical clustering of the ISPPs was performed using complete linkage, using Euclidean distance on the p-values derived from multiple comparison. No scaling or transformation was applied to the p-values. The visualization of the data was facilitated by ClustVis (36) (https://biit.cs.ut.ee/clustvis/).

In contrast, Categories B, C, and D exhibited tight clustering (Figure 4A). Within Category B, FATHMM (20) was anomalously distant, correlating with its high p-values (see Data Availability) and reduced number of significant genes (Figures 2, 3). Similarly, EVE (38) from Category D deviated significantly from its cluster, performing less effectively across the exome and Mendeliome (Supplementary Figure 2, Figure 3), despite integrating multiple parameters.

ISPPs in Category C demonstrated tight clustering for the Mendeliome (Figure 4A), reflecting consistent predictions and focus on biological function within a dataset of disease-causing genes.

Hierarchical clustering (Figure 4B) confirmed the relevance of the initial ISPP categorization (Figure 2), with ISPPs in the same category generally clustering together. Exceptions were observed for three ISPP pairs from different categories: phyloP100way-V (16) and fathmm-MKL (39), DEOGEN2 (40), and PROVEAN (21), and VEST4 (22) and BayesDel_noAF (35), indicating occasional overlaps in categorical boundaries, but always with the adjacent.

It was shown that ISPPs from the same algorithmic approaches are clustered together for Benign variant assessment (Figure 5A). In contrast, ISPP clusters for pathogenic variants are broader and involve ISPPs from multiple categories (Figure 5B). For both variant groups, most ISPPs within the same category also tended to cluster closely, as FATHMM (20) showed the most divergent results, consistent with previous findings (Figure 4).

**Figure 5.**
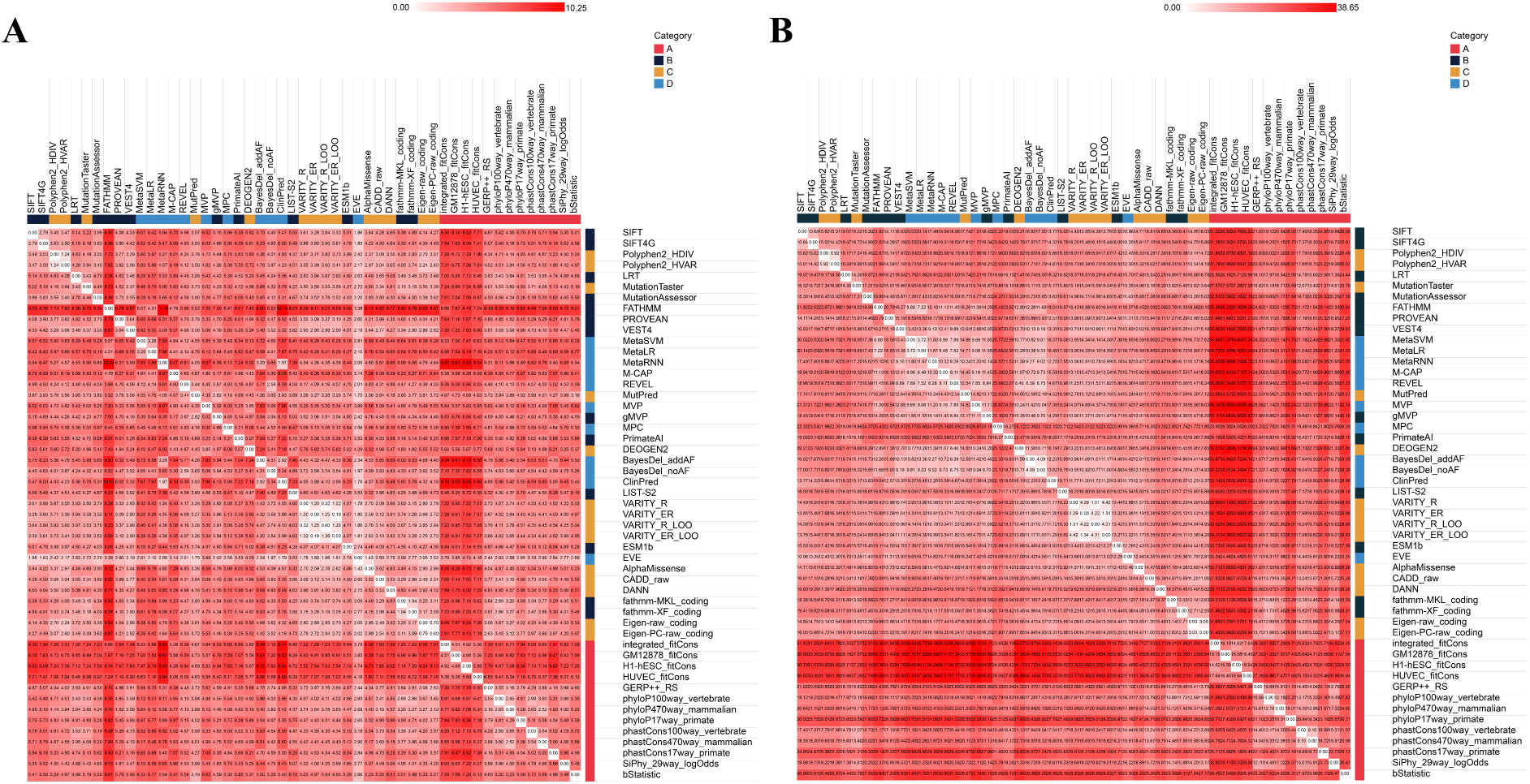
ISPP similarity matrix for. **A.** Benign and **B.** Pathogenic variant assessment. This heatmap uses the Euclidean distance metric to illustrate similarity among ISPPs for the variants based on ISPP scores. Darker red shades indicate greater distances. ISPP categories are color-coded as indicated in the legend.

Despite its divergence, FATHMM (20), in benign variant group, shared similarities with ISPPs from categories C and D (e.g. MetaSVM (41), MetaLR (41), M-CAP (42), REVEL (43), MVP (44), EVE (38), MutPred (45)) highlighting a closer proximity between FATHMM and these categories (Figure 4) for Benign variant assessment.

fitCons-derived ISPPs (37) maintained tight internal clustering but diverged from category A and other ISPPs, aligning with earlier results (Figure 3, Figure 4). Conversely, algorithms like phyloP (16) and phastCons (17) showed greater similarity in benign variant scoring both within and across other ISPPs, underscoring their consistency in benign variant evaluation (Figure 4). In pathogenic variant assessment, MPC (46) and PrimateAI (47) diverged markedly from their categories and other ISPPs.

### 2.4 VAMPP-score

All variants from dbNSFP v4.7A (1, 2) were annotated with VAMPP-score. A total of 23,464,253 variants had available results.

The benign variant distribution (Figure 6) is characterized by a concentration of lower scores, with a peak approximately at 0.1, while the distribution for pathogenic variants is broader, peaking around 0.3. Although both distributions exhibit a unimodal pattern with a single peak, the pathogenic group’s distribution is less symmetrical than that of the benign variants. The tail of the pathogenic distribution extends toward higher VAMPP-scores, suggesting an effective differentiation of pathogenic from benign variants.

**Figure 6.**
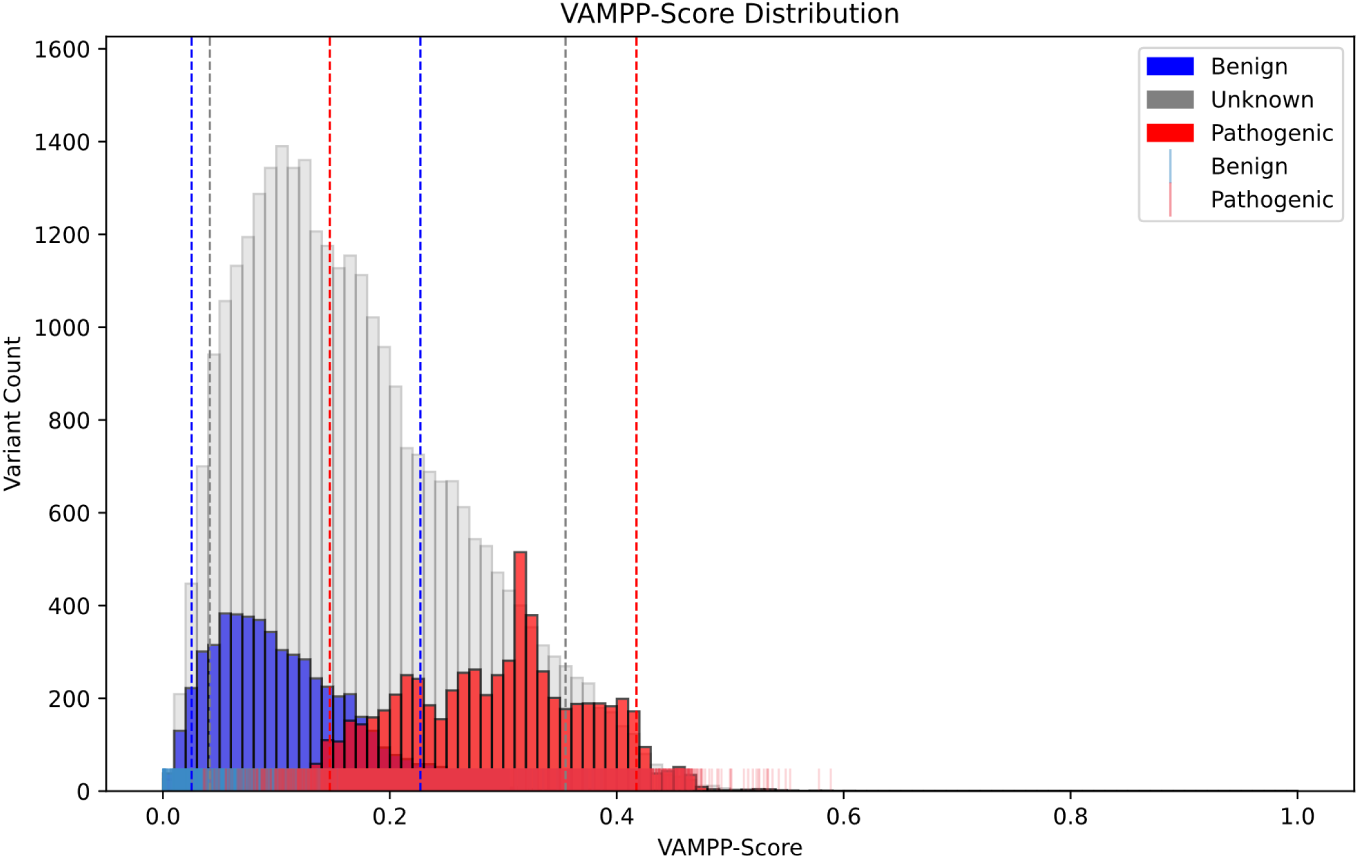
The histogram illustrates the VAMPP-score distribution within the human exome, categorizing variants as Benign, Unknown, and Pathogenic. The x-axis denotes the VAMPP-scores, while the y-axis indicates the count of variants within each group. Beneath the histogram, a rug plot visualizes individual variant scores for the Benign and Pathogenic groups. Quantile lines (dashed lines according to the color legend) are marked for each group.

The two variant distributions overlap in the range of approximately 0.1-0.3. Out of 3,245 pathogenic and 2,585 benign variants that are in the overlap range, 1,901 of the pathogenic and 1,964 variants benign group were under the review status of criteria provided, single submitter. This suggests that review status and submitter categories in ClinVar (3) can be considered as an indicator for possible false negative/positive results. A variant that has been submitted as pathogenic multiple times without conflicting interpretations (ClinVar-two stars) is more reliable and supported by the literature than a variant that has been submitted only once or with conflicting interpretations (ClinVar – one star).

#### 2.4.1 Variant Evaluation with VAMPP-score

The VAMPP-score is suggested for use in variant interpretation. In this context, we defined cut-offs for ACMG criteria (11) in multiple lines of computational evidence (PP3 and BP4) and evidence-level adjustments for variant assessment. Since a distinction between the known P, LP, P/LP, and B, BL, B/LB variants was observed by VAMPP-score, we incorporated the quantile information and the distributions of the variants with known clinical significance to this cut-off determination (see Section 4.2 for details). Benign variants show quantiles at 0.0252 for the 5th percentile and 0.226 for the 95th percentile, Unknown variants at 0.041 for the 5th percentile and 0.354 for the 95th percentile, and Pathogenic variants at 0.147 for the 5th percentile and 0.41 for the 95th percentile (Figure 6).

As a result, we defined a set of thresholds for ACMG-PP3/BP4 criteria adjustments:

● A cut-off value of 0.226 was determined for a variant to be considered as benign, and the scores less or equal to that would indicate benignity. In addition, the pathogenicity cut-off value was determined as 0.354, where variants with scores higher or equal to would be considered pathogenic. In both cases, we suggest that multiple in-silico evidence criteria strength level may be upgraded to moderate and strong for both PP3 and BP4, respectively.

VAMPP-score’s performance analysis is under making within the ISPPs available in dbNSFP4.7A (1, 2).

## 3 Discussion

In variant interpretation, various ISPPs provide insights on clinical significance using publicly available data, which, despite ensuring accurate predictions, may generalize results across populations and genetic events. The limitations and biases in these datasets must be recognized, as experts often rely on ISPPs based on habits and experience. Previously, the prediction performances and accuracies of the ISPPs have been investigated in several studies (10, 48–50), and a combinatorial approach has been suggested for the classification of Rett syndrome-associated gene variants (51). Here we aimed to analyze the prediction performances of the ISPPs, creating a statistical framework with proximity functions, and developed the VAMPP-score, available soon at www.vamppscore.com.)

We utilized data from dbNSFP (1,2), including 52 ISPPs for missense variants interpretation, categorizing them into four based on their methodologies (Figure 2). Liu et al. (2) previously analyzed the distribution and inter-correlations of ISPPs in 2020. Among the 45 ISPPs analyzed, fitCons (37) scores showed weak correlations with other scores but were internally consistent. Similarly, bStatistic (52) scores exhibited weak correlations with other scores. Our analysis indicated that fitCons had the lowest performance across the exome and Mendeliome, followed by bStatistic, as evidenced by clustering and similarity analyses (Figure 3, Figure 4, and Figure 5). Liu et al. also compared the agreement on variant predictions for 20 ISPPs in dbNSFP, noting that highly agreed ISPPs, such as BayesDel (35) and PolyPhen (53) pairs, were trained on the same data, consistent with our findings.

Our results revealed clustering of categorized ISPPs, e.g. C and D, corresponding to ensemble prediction scores in dbNSFP (1, 2). Unlike Liu et al. (2), who used Pearson correlation coefficients on rank scores, we applied Euclidean distance with complete linkage among p-values from multiple comparison tests, representing each ISPP’s performance in distinguishing variant groups. Our supervised approach involving novel ISPP categorization (Figure 2) confirmed similar prediction performances of ISPPs in the same category for known pathogenic and benign variants (Figures 4 and 5). The resulting clusters demonstrated that ISPPs categorized as meta-predictors can be subdivided (Figures 2 and 4), emphasizing the robustness of our categorization method. Liu et al. also evaluated the performance of 45 ISPPs using ClinVar (3) and gnomAD (28) data, identifying top scores for different groups. Our findings align, showing top ISPPs mostly grouped in category D, including BayesDel_addAF (35), ClinPred (27), REVEL (43), and MetaLR (41), with MetaRNN (34) identified as the best predictor for both exome and Mendeliome (Figure 3), corroborating its superior performance in our analysis.

Category A ISPPs had the fewest significant genes, possibly due to limitations in nucleotide-based predictions (Figure 3). These ISPPs utilize multiple sequence alignments to identify conserved genomic regions at the nucleotide level. This can be misleading for missense variants. The proteins are composed of amino acids, most of which can be coded by multiple codons (54, 55). This redundancy means nucleotide changes do not always result in amino acid changes. Consequently, ISPPs in Category A might inaccurately predict missense variant effects where nucleotide conservation is low, but protein sequence conservation is maintained. Despite these limitations, ISPPs remain crucial for evaluating evolutionary conservation in non-coding regions fitCons (37) was found to have the least performance on accurately predicting missense variant pathogenicity. It is an algorithm that relies on fitness consequences that can be interpreted as an evolution-based measure of potential genomic function. It was mainly raised to solve the problem of identifying non-coding functional elements. Even though the scores were also considered to be elevated for coding regions in the genome it is described as considerably better sensitivity for the noncoding regions. This correlates with our results and suggests that for a more accurate performance evaluation, fitCons scores should be investigated, focusing on the noncoding regions of the human genome.

The primary data source used is ClinVar (3), which has an uneven gene-variant distribution. This disparity indicates that some genes are more studied, and hence more data is submitted because they are more *popular* and chosen for further studies (56). Consequently, insufficient variant data was available for certain genes in our analysis. From ClinVar’s limited data, only 23,464,253 out of over 84 million variants in dbNSFP v4.7A (1, 2) were annotated with VAMPP-scores for 3,955 genes. This is partly because benign variants, lacking clinical significance, are often underreported compared to pathogenic variants. Benign variants are usually dismissed due to their high population allele frequencies and polymorphisms (57–59). Additionally, severe variants may not be observed due to their incompatibility with life, often being eliminated through evolution or causing early death, leading to their underrepresentation in clinical databases. Moreover, variants might not be observed in the population due to severe consequences caused by their incompatibility with life. Variants occurring in the conserved regions of the genome, causing severe effects and dysfunctions, may be eliminated through evolution (18, 60). Also, variations occurring in these regions may cause prenatal or early death (61), leading to their unrecognition.

The VAMPP-score distribution shows a concentration within a specific range (Figure 6), not indicating inaccuracies but highlighting the potential for misleading interpretations when ISPP scores are viewed individually. Evaluating variant pathogenicity based on ISPP rank scores is good practice; however, ISPPs can have varying score distributions. One ISPP may have scores near 1, while another may have lower scores for known pathogenic variants, complicating interpretation and potentially leading to inconsistent conclusions. The VAMPP-score’s proximity estimation function and mean difference help mitigate issues from these unequal distributions. The broad range of VAMPP-scores suggests significant differences in variant characteristics, distinguishing between P/LP and B/LB groups (Figure 6). Pathogenic variants generally have higher scores, showing that the VAMPP-score effectively integrates gene-ISPP pairs. However, uncertainty remains for intermediate-scoring variants, often classified as Variants of Uncertain Significance (VUS) in ClinVar (3) reviews, depending on clinical contexts.

The ISPPs’ prediction performance for gene counts across the exome and Mendeliome revealed a consistent top-eight order (Figure 3, Supplementary Figure 2). Intriguingly, five of these top ISPPs (MetaRNN (34), ClinPred (27), BayesDel_addAF, BayesDel_noAF (35), MetaLR(41)) belong to category D, indicating the high accuracy of ensemble predictors This aligns with VarSome’s (62), findings, which recently implemented an approach for ACMG-PP3 criterion evaluation (63). MetaRNN (145) was shown as the most accurate ISPP, showing 91% accuracy in classifying ClinVar (3) variants, consistent with our results. The rest of the top five ISPPs, except ClinPred (27), also matched our accuracy patterns.

The ACMG/AMP 2015 guidelines for variant interpretation (11) are essential for clinicians and genetic analysts. These guidelines offer a scoring metric to aid interpretation but do not replace clinical judgment. Clinicians must combine these metrics with a comprehensive evaluation, considering the patient’s phenotype, gene inheritance patterns, clinical manifestations of the variant, and potential genetic phenomena. The guidelines specify criteria for computational evidence on pathogenicity (PP3) and benignity (BP4), requiring agreement among ISPPs and serving as supporting evidence. Previously, studies have shown that these criteria can be adjusted when specific thresholds are met for the ISPPs individually (10, 48, 64). Our approach integrates ISPP performances into a combinatory statistical framework, resulting in several recommendations: *(i)* Using the top eight ISPPs revealed here for general missense variant interpretation practices for Mendeliome, *(ii)* Including the result of one ISPP from each category from this top 8 when interpreting a variant and deciding on the final assessment upon their agreement, *(iii)* Including the results from the best-performing ISPPs for the gene where the variant of interest is located for a more conserved evaluation, which is one of the components of the VAMPP-score used in weighting.

The VAMPP-score, with its mathematical foundation, highlights the best-performing ISPPs for each gene and integrates their performance into the final score. Adjusting ACMG criteria levels using VAMPP-score cut-offs can significantly influence variant classification. For instance, upgrading the ACMG-PP3 criterion to moderate for pathogenicity if a variant’s VAMPP-score meets the conservative pathogenic threshold, and similarly defining the BP4 criterion as strong for benign interpretations. If the variant’s VAMPP-score meets the second set of thresholds, the PP3/BP4 criterion can be included in the evaluation, preferably with following the recommendations proposed in this study for ISPP interpretations. Reevaluation of variants using VAMPP-score can lead to reclassification of variants previously considered VUS to likely pathogenic (LP). This reclassification can facilitate definitive genetic diagnoses and targeted treatments, particularly in rare disorders. Despite its utility, some variants may still be classified as VUS due to limited data or evidence (10), suggesting functional studies. However, the VAMPP-score can be used to prioritize candidate variants for such analyses.

The novel approach described here offers new insights into computational methods. By using statistical methods to compare ISPPs and incorporating the best-performing gene-based ISPPs, we can reclassify previously VUS variants with diagnostic value. Our integration of key components essential for detecting pathogenicity has led to the development of the VAMPP-score. This adaptable framework can also be applied to other variant data types, like somatic variations and noncoding regions, given available curated data. By following our variant interpretation strategies and using the VAMPP-score, we aim to enhance the credibility and acceptance of computational methods, reducing existing mistrust and biases. Ultimately, we believe that variant classifications supported by computational methods and clinical outcomes will lead to more positive results in clinical genetics and diagnosis.

We are actively validating the VAMPP-score using clinical cases that have been diagnosed primarily through clinical expertise. Many of these cases involve Variants of Uncertain Significance (VUS), and our goal is to determine if we can reclassify them without the need for further time-consuming analyses, such as segregation analysis. This ongoing validation process aims to prioritize these variants for genetic diagnosis, ensuring the robustness and practical applicability of our framework in real-world clinical genetics. Through comprehensive analysis of clinical data, we seek to demonstrate the VAMPP-score’s potential to enhance genetic diagnostics and improve patient outcomes. As we continue to gather and analyze more clinical data, we are committed to refining and optimizing the VAMPP-score to ensure its effectiveness and credibility in the field of genetic research.

## 4 Materials and Methods

### 4.1 Data curation and database structure

#### 4.1.1 The dbNSFP database

All possible single nucleotide variations in the human genome, over 84 million variants, with their corresponding information provided by dbNSFP4.7A (1, 2) was used to create a DuckDB (65) (version 0.9.3-dev) database (Supplementary Figure 6), which allows fast and efficient querying and extraction of the initial dataset based on specified criteria of choice for downstream analysis.

The utilization of DuckDB (65) was preferred, especially because of the need for an updatable database. The datasets integrated into the structure are composed of genomic variation data (Supplementary Figure 6), at least one of which is known to get updated weekly (3). In addition, the dbNSFP (1, 2) data has been getting updates more frequently for the past two years (66), due to advancements in the field, development of new computational predictors, updates, and error fixes.

#### 4.1.2 Canonical transcript mapping

Multiple sequence ontology annotations for variants chosen to be included in the analysis were aimed to be avoided. For this, canonical transcript IDs for each gene based on Ensembl Release 111 (January 2024) (29) were integrated into our DuckDB (65) database as a new table. During query extraction, variants that have matching transcript IDs within dbNSFP (1, 2) and the reference variant list in pathogenicity assessment from ClinVar (151) were chosen upon Ensembl canonical transcripts (Supplementary Figure 6).

#### 4.1.3 ClinVar variants

The updated ClinVar (3) variant summary data was incorporated as a reference variant list for pathogenicity assessment (March 2024). The variant summary data was accessed through the ClinVar FTP site, and it was pre-filtered based on the following criteria: missense variants, annotated according to the genome assembly GRCh38 annotations with unspecified review status. The variants were further refined upon their clinical significance to avoid the involvement of conflicting interpretations and to ensure that more evident clinical outcomes were included in the analysis. Accordingly, the variants with the following clinical submissions were kept: Pathogenic (P), Likely pathogenic (LP), Pathogenic/Likely pathogenic (P/LP), Uncertain significance (VUS), Likely benign (LB), Benign/Likely benign (B/LB), and Benign (B).

It was strategically chosen to utilize the ClinVar variant list without imposing restrictions on the review status of the variants, a decision primarily influenced by the need to increase the statistical power of the analysis. Since the restriction of the variants according to their review status significantly reduces the number of variants in the list, this inclusive approach of no restrictions allows for a more comprehensive examination of the variants with known clinical significance with at least a submission level to a database.

#### 4.1.4 Final dataset for downstream analysis

VAMPP-score utilizes the best gene-ISPP matches by integrating several parameters into the calculation, including the results from statistical tests and proximity metrics. A final dataset to calculate metrics and perform analyses (ClinVar_variant_summary_March24) was generated based on canonical transcript ID and refined ClinVar (3) variant data – transcript ID-match status, querying a list of variants with the necessary information available in dbNSFP (1, 2).

Due to the size of the dbNSFP dataset, only specific columns, including general information on the variants and ISPP scores and rank scores, were extracted to avoid irrelevant information handling and computational exceeding.

### 4.2 VAMPP-score

As a part of the VAMPP-score, a statistical framework of multiple and pairwise comparisons was created. Firstly, for multiple comparisons, Kruskal-Wallis test (30)was used for each gene-ISPP pair to determine if there is a significant pathogenicity score difference (p<0.05) among the three independent variant groups, Pathogenic, Benign, and Unknown. Subsequently, the Wilcoxon-Rank-Sum test (67) was applied for each gene-ISPP pair to detect a significant difference in scoring (p<0.05) among variant groups. These groups were “Pathogenic vs. Benign”, “Pathogenic vs. Unknown”, and “Unknown vs. Benign”. By utilizing Wilcoxon-Rank-Sum, it was aimed to determine how the variant groups differ from each other and/or whether or not they come from the same distribution.

The ISPPs with p<0.05 were integrated into the VAMPP-score as a part of the weighted gene coefficient calculated for each gene-ISPP pair. In this context, the ISPPs with p-values less than 0.05 from multiple comparisons and ISPPs with *p<0.05* from Pathogenic vs. Benign pairwise comparisons were included in the calculation with their p-values subtracted from 1 to reflect the power of the ISPP’s significance.

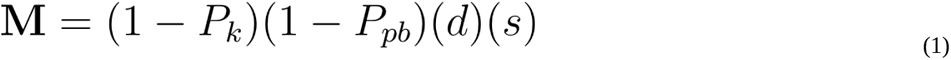

The proximity (*d*) estimation function (PEF) (Formula 1.1) is based on the assumption of how the three predefined variant groups should be distributed by means of assigned pathogenicity scores by each ISPP. In this context, the means of each variant group for each gene-ISPP pair were calculated and integrated with the VAMPP-score, considering their ideal distribution (Figure 8).

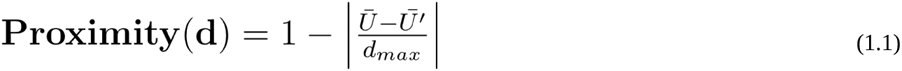

**Figure 8.**
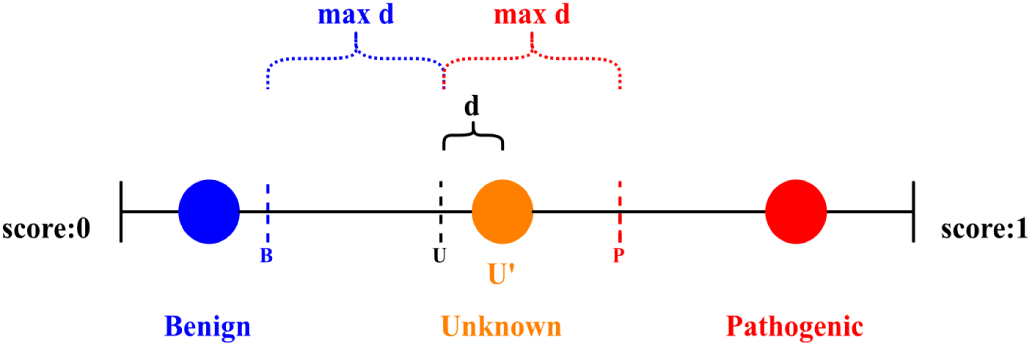
Ideal variant group distribution. The figure shows the ideal variant group distribution for a gene based on the assigned pathogenicity scores from each ISPP. Filled circles represent the ideal means of the defined variant groups as Benign (blue), Unknown (orange), and Pathogenic (red). Dashed lines represent any variant group mean for Benign (B), Unknown (U), and Pathogenic (P) for an ISPP, respectively. “max *d*” represents the maximum distance, which is the numerical expression of how much the mean of the Unknown variant group diverges from and serves as an indicator of how it is positioned among the other two variant groups for each gene-ISPP pair.

To calculate *d*, a theoretical Unknown variant group mean (U’) is defined as the median of pathogenic and benign variant group means for each gene-ISPP pair. Then, relying on the assumption that U’ (Formula 1.1.1) would be equidistant from the Pathogenic or Benign variant groups,

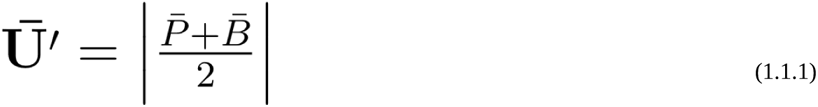

the maximum distance *d_max_* (Formula 1.1.2) is calculated by subtracting the U’ from the Pathogenic variant group mean (Figure 8).

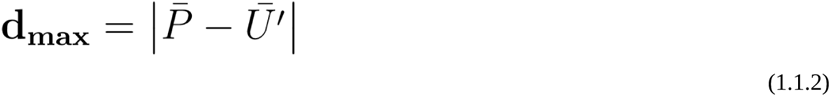

Finally, U’ is subtracted from the actual mean of the unknown variant group (U) for each gene-ISPP pair and divided by *d_max_*. The absolute resulting value is subtracted from 1 to be referenced as a significant prediction accuracy value and included in the final VAMPP-score calculation (Formula 1) as *d* (Formula 1.1).

A score difference multiplier is another element of the VAMPP-score, the final element included in M, described as *s* (Formula 1.2). *s* ultimately determines the sign of M for each gene-ISPP pair by checking whether the Pathogenic variants were scored higher than Benign considering the rank score range of 0-to-1.

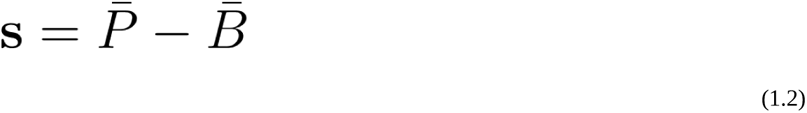

After calculating all the parametric values of M, the VAMPP-score was calculated (Formula 2) for each variant available in dbNSFP v4.7A (1, 2), then scaled between 0 and 1 by stretching the values as the minimum value was made equal to 0, and the maximum value became 1 using a linear normalization for ease of comparison.

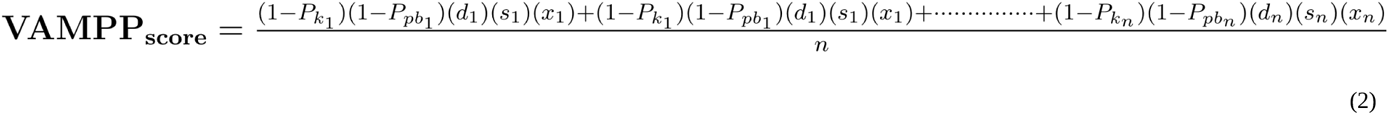

In VAMPP-score, *M* stands for the gene of interest where the variant is located, is used as an impactor, and is multiplied by the rank score of the ISPP of interest (x _1_, x_2_,..x_n_) for that variant. Addition is performed for the number of ISPPs where there is an available *M* for the gene-ISPP pair, and the sum is then divided by the number of the ISPPs included in the calculation for the gene where the variant of interest is located.

After variant annotation with VAMPP-score, to define cut-offs for pathogenicity and benignity, quantile lines and the distribution of the scores within each group were analyzed. Threshold values to define pathogenicity and benignity were identified:

To classify a variant as benign, the VAMPP-score should be less than the 95th percentile of benign variants and lower than the 5th percentile of unknown variants. This can be set around as the more conservative approach to use 0.226 as the upper limit to avoid overlap with unknown variants. On the other hand, to classify a variant as pathogenic, the VAMPP-score should be greater than the 95th percentile of benign variants and higher than the 5th percentile of pathogenic variants. It could be set around 0.354 to avoid overlapping with unknown variants.

### 4.3 Multidimensional clustering and correlation analysis

The results obtained from the multiple comparison tests were also used to determine gene-based ISPP consistency. In this context, for multidimensional clustering, principal component analysis (PCA) (68, 69) was performed for categorized ISPPs utilizing ClustVis v1.0 (36). PCA helps to reduce the dimensionality of the data with a large set of variables, hence giving a generalized representation of the data by still keeping all the variables. PCA was used to cluster ISPPs on their significance, and the unit variance scaling method was applied to rows (Mendeliome genes) for scaling, along with SVD with imputation as the PCA method. The ISPP performance on the Mendeliome gene list was also visualized as a heatmap ClustVis v1.0, no scaling and transformation were applied to the initial data, and raw p-values were observed in the original 0-1 scale.

Hierarchical clustering of the ISPPs was performed using complete linkage utilizing Euclidean distance (180), where clusters were merged based on the maximum distance between points in the clusters. ISPP performance and their variant-based pathogenicity prediction score correlation analyses were carried out. Three different similarity matrices for ISPPs were created and visualized utilizing Broad Institute’s Morpheus (https://software.broadinstitute.org/morpheus): a similarity matrix created with ISPP p-values from multiple comparisons (Supplementary Figure 4), and two other similarity matrices created with the ISPP rank scores for Benign (Figure 5) and Pathogenic (Figure 6) group variants. The Euclidean distance metric was used in similarity matrices to detect the similarity between ISPPs upon their p-values from multiple comparisons and within Pathogenic and Benign variant groups upon rank scores from ISPPs.

The general distribution of the VAMPP-score throughout the genome was visualized using a histogram by Python v.3.10.12’s Matplotlib (version 3.8.4) (70). In addition, VAMPP-score distribution for Pathogenic, Benign, and Unknown variant groups was plotted as a histogram, showing the variant count for each group utilizing Matplotlib. Its distribution was combined with a rug plot to represent individual data points for Pathogenic and Benign variants using the Matplotlib library in Python (version 3.10.12).

## Data Availability

The datasets generated during and/or analyzed during the study are available from the corresponding author upon reasonable request. Also, the annotation data of VAMPP-score, gene scores representing the best-performing ISPPs per gene, and statistical data are available through our GitHub page (https://github.com/eylulaydin7/VAMPP-score).

