## Supplemental Data for "A New Era in Missense Variant Analysis: Statistical Insights and the Introduction of VAMPP-Score for Pathogenicity Assessment"

**Supplementary Table 1.** In-silico pathogenicity predictors (ISPP) table. The table shows the most frequently used ISPPs with rank scores available in the latest version of dbNSFP v4.7A (1, 2). Definitions of ISPP acronyms were written below the name of the ISPP, if available.

| Predictor | Subtype, if any | Basic information | Reference |
|---|---|---|---|
| **GERP**<br>**G**enomic **E**volutionary **R**ate **P**rofiling | _RS: rejected substitution | identifies evolutionary constrained elements in multiple-species alignments by comparing rates of nucleotide substitutions | (18) |
| **phyloP** | 17way: primate species list<br>100way: vertebrate species list<br>470way: mammalian species list | measures evolutionary conservation at individual alignment sites by assessing the likelihood of observing the given number of substitutions under a neutral model | (16) |
| **phastCons** | 17way: primate species list<br>100way: vertebrate species list<br>470way: mammalian species list | estimates the probability of each nucleotide being conserved based on a phylogenetic hidden Markov model | (17) |
| **SiPhy** | 29way: mammalian species list | utilizes patterns of nucleotide substitution and insertion/deletion events to identify regions under evolutionary constraint | (71) |
| **bStatistic** | | quantifies the degree of background selection at a given site in the human genome | (52) |
| **gMVP** | | a genomic variant pathogenicity prediction tool that integrates multiple genomic features on structure of graph neural network, a version of MVP | (72) |
| **LIST-S2**<br>**L**ocal **I**dentity and **S**hared **T**axa-**S2** | | assesses local sequence conservation and shared taxa to predict pathogenicity | (73) |
| **FATHMM**<br>**F**unctional **A**nalysis **t**hrough **H**idden **M**arkov **M**odels | _MKL: multiple kernel learning (39)<br>_XF: eXtended features (74) | predicts the functional effects of missense variants using hidden Markov models | (20) |

**Supplementary Table 1.** In-silico pathogenicity predictors (ISPP) table. The table shows the most frequently used ISPPs with rank scores available in the latest version of dbNSFP v4.7A (1, 2). Definitions of ISPP acronyms were written below the name of the ISPP, if available (continued).

| Predictor | Subtype, if any | Basic information | Reference |
|---|---|---|---|
| **SIFT**<br>**S**orting **I**ntolerant **f**rom **T**olerant | _4G: for genomes | predicts whether an amino acid substitution affects protein function based on sequence homology and the physical properties of amino acids | (19) |
| **EVE**<br>**E**volutionary Model of **V**ariant **E**ffect | | uses a machine learning approach to predict the functional impact of coding variants based on evolutionary information | (38) |
| **LRT**<br>**L**ikelihood **R**atio **T**est | | assesses whether a nonsynonymous (amino acid changing) variant is likely to be pathogenic based on a likelihood ratio test | (75) |
| **VARITY** | _R:rare<br>_ER:extremely rare<br>_R/ER-LOO:leave-one-out | predicts the impact of amino acid substitutions on protein function, has subtypes on rare and extremely rare variants, with LOO approach | (76) |
| **MutationAssessor** | | predicts the functional impact of amino acid substitutions in proteins based on evolutionary conservation | (77) |
| **MutationTaster2** | | predicts disease-causing potential of genetic variants including SNPs and indels | (78) |
| **VEST4**<br>**V**ariant **E**ffect **S**coring **T**ool | | uses a machine learning approach to score the pathogenicity of missense mutations | (22) |
| **PROVEAN**<br>**Pro**tein **V**ariation **E**ffect **An**alyzer | | predicts whether a protein sequence variation is likely to be deleterious | (21) |
| **MutPred** | | predicts the potential pathogenic effects of AA substitutions in humans | (45) |
| **PrimateAI** | | predicts disease-causing variants based on a deep learning model trained on primate data; the human+primate deep learning network | (47) |

**Supplementary Table 1.** In-silico pathogenicity predictors (ISPP) table. The table shows the most frequently used ISPPs with rank scores available in the latest version of dbNSFP v4.7A (1, 2). Definitions of ISPP acronyms were written below the name of the ISPP, if available (continued).

| Predictor | Subtype, if any | Basic information | Reference |
|---|---|---|---|
| **DEOGEN2** | | incorporates heterogeneous information about the molecular effects of the variants, the protein domains, and gene interaction information | (40) |
| **AlphaMissense** | | makes variant effect prediction proteome-wide, focuses on missense variants; a deep learning model that builds on AlphaFold2 for protein structure predictions | (25) |
| **PolyPhen2**<br>**Poly**morphism<br>**Phen**otyping v**2** | _HDIV & _HVAR: two data sets | makes predictions for AA substitutions on the protein structure and function using physical and comparative considerations | (53) |
| **fitCons**<br>**fit**ness **Cons**equences | integrated_fitCons<br>GM12878(human lymphoblastoid cells) fitCons<br>H1-hESC(human embryonic stem cells)_fitCons<br>HUVEC(human umbilical vein epithelial cells)_fitCons | assesses the selective constraint on individual nucleotides within genomic regions | (37) |
| **MVP**<br>**M**issense **V**ariant **P**athogenicity prediction | | prediction method that uses deep residual network to predict the effect of missense variants | (44) |
| **BayesDel** | _noAF: no allele frequency information added<br>_addAF: allele frequency information added | combines multiple predictors using a Bayesian framework, utilizes the approach PERCH (Polymorphism Evaluation, Ranking, and Classification for a Heritable trait) | (35) |
| **CADD**<br>**C**ombine **A**nnotation **D**ependent **D**epletion | _19: considers hg19 as the reference genome | integrates multiple annotations into a single metric to predict the deleteriousness | (23) |

**Supplementary Table 1.** In-silico pathogenicity predictors (ISPP) table. The table shows the most frequently used ISPPs with rank scores available in the latest version of dbNSFP v4.7A (1, 2). Definitions of ISPP acronyms were written below the name of the ISPP, if available (continued).

| Predictor | Subtype, if any | Basic information | Reference |
|---|---|---|---|
| **DANN** | | a deep learning approach that utilizes a neural network for prediction, trained on CADD data | (24) |
| **Eigen_coding** | _PC: principal component, uses the lead eigenvector for weighting annotations | uses a machine learning model to predict the functional effect of coding variant | (79) |
| **GenoCanyon** | | statistical method to predict functional regions in the human genome, can be used for both coding and non-coding regions | (80) |
| **ClinPred** | | predicts the pathogenicity of missense variants based on clinical data | (27) |
| **MetaSVM** | | support vector machine-based method integrating multiple predictors to assess variant pathogenicity. | (41) |
| **MetaLR** | | uses logistic regression to integrate multiple variant effect predictors. | (41) |
| **MetaRNN** | | recurrent neural network approach integrating various predictors | (34) |
| **M-CAP**<br>**M**endelian **C**linically **A**pplicable **P**athogenicity | | predicts the pathogenicity of missense variants in Mendelian disorders | (42) |
| **REVEL**<br>**R**are **E**xome **V**ariant **E**nsemble **L**earner | | ensemble method combining multiple tools to predict the pathogenicity of rare missense variants | (43) |
| **MPC**<br>**M**issense badness, **P**olyPhen-2, **C**onstraint | | combines missense deleteriousness, PolyPhen-2, and constraint scores | (46) |
| **ESM1b**<br>**E**volutionary **S**cale **M**odeling | | uses evolutionary modeling to predict the effects of protein mutations | (81) |
| **LINSIGHT**<br>**L**inear **I**nference of **N**atural **S**election from **I**nterspersed **G**enomically co**H**erent elemen**T**s | | predicts functional significance of genomic variants based on evolutionary conservation | (82) |

**Supplementary Table 2. The details for the input ClinVar (3) variants used in the analysis for Exome and Mendeliome.** (P: Pathogenic, LP: Likely pathogenic, P/LP: Pathogenic/Likely pathogenic, B: Benign, LB: Likely benign, B/LB: Benign/Likely benign)

| | Genes | Total Variants | Pathogenic | Benign | Unknown |
|---|---|---|---|---|---|
| ***Exome*** | 17,712 | 1,162,673 | 57,741 | 97,740 | 1,010,462 |
| | | | P: 23,077 | B: 28,875 | |
| | | | LP: 26,206 | LB: 55,273 | |
| | | | P/LP: 8,188 | B/LB: 10,592 | |
| ***Mendeliome*** | 4,703 | 841,674 | 55,253 | 67,347 | 719,074 |
| | | | P: 22,236 | B: 20,340 | |
| | | | LP: 25,085 | LB: 37,099 | |
| | | | P/LP: 7,932 | B/LB: 9,908 | |

**Supplementary Table 3.** Gene counts for ISPPs. The table shows the ISPPs and the number of genes with significant results (p-value <0.05) for that ISPP throughout Mendeliome. (The ISPPs were listed according to the gene counts descendingly.)

| ISPP | Gene count |
|---|---|
| MetaRNN | 3027 |
| ClinPred | 2960 |
| Bayesdel_addAF | 2895 |
| VEST4 | 2441 |
| AlphaMissense | 2400 |
| Bayesdel_noAF | 2368 |
| MetaLR | 2279 |
| VARITY_R | 2264 |
| REVEL | 2222 |
| VARITY_R_LOO | 2222 |
| VARITY_ER | 2214 |
| VARITY_ER_LOO | 2203 |
| gMVP | 2155 |
| CADD | 2130 |
| Polyphen2-HVAR | 2048 |
| M-CAP | 2030 |
| Eigen-raw_coding | 2025 |
| DEOGEN2 | 2023 |
| ESM1b | 2017 |
| SIFT4G | 2004 |
| SIFT | 1988 |
| Eigen-pc-raw_coding | 1971 |
| PROVEAN | 1970 |
| Polyphen2-HDIV | 1930 |
| MutationAssessor | 1924 |
| MutPred | 1909 |
| MetaSVM | 1906 |
| MPC | 1900 |
| PrimateAI | 1874 |
| LIST-S2 | 1796 |
| fathmm-XF_coding | 1794 |
| MVP | 1742 |
| MutationTaster | 1720 |
| fathmm-MKL_coding | 1689 |
| phyloP100way_vertebrate | 1647 |
| DANN | 1549 |
| LRT | 1474 |
| phyloP470way_mammalian | 1371 |
| phastCons100way_vertebrate | 1366 |
| phastCons470way_mammalian | 1280 |
| GERP++_RS | 1247 |
| SiPhy_29way_logodds | 1214 |
| FATHMM | 1183 |
| EVE | 1073 |
| phastCons17way_primate | 862 |
| phyloP17way_primate | 853 |
| bStatistic | 675 |
| h1-hesc_fitcons | 569 |
| GM12878_fitcons | 567 |
| HUVEC_fitcons | 558 |
| integrated_fitcons | 546 |

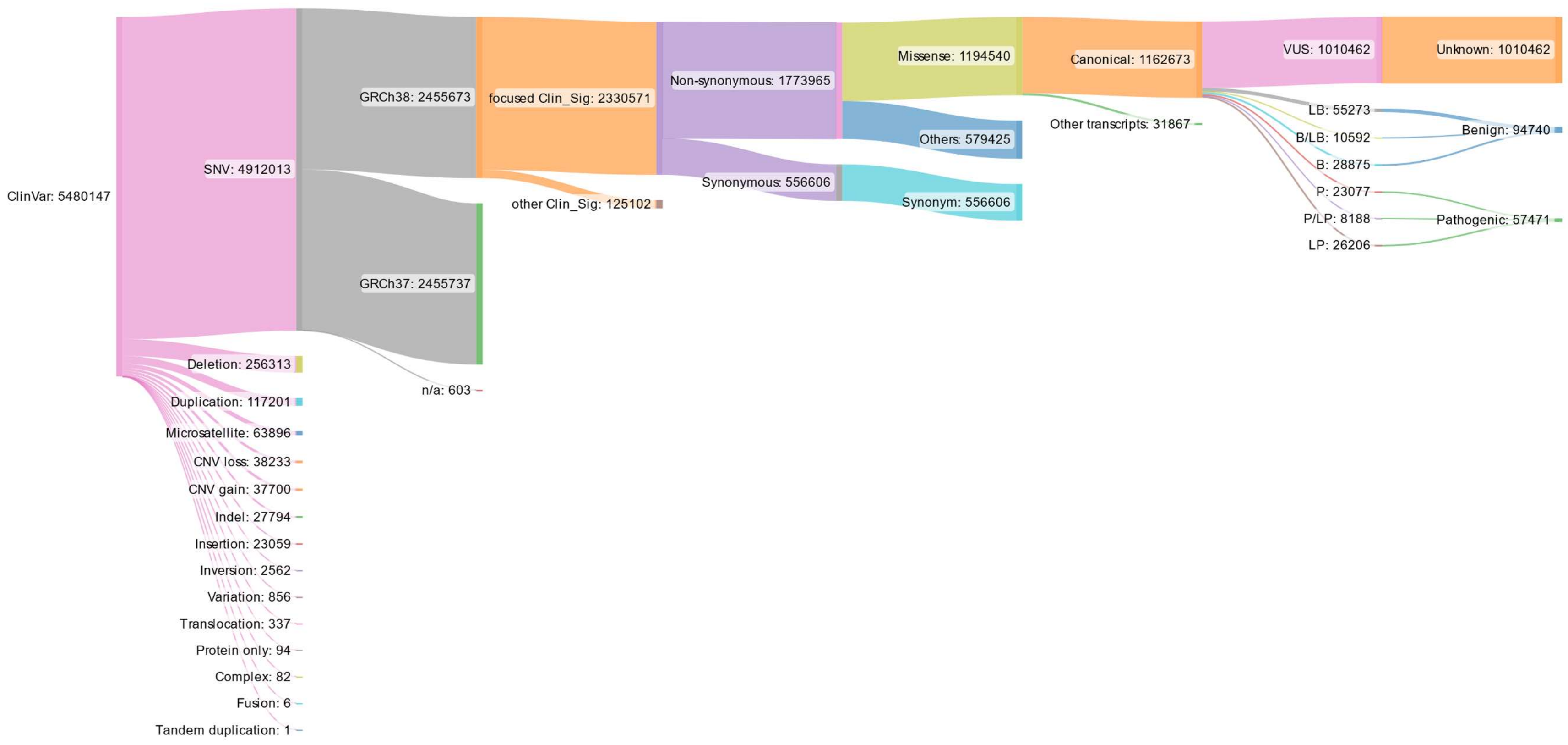

**Supplementary Figure 1. Schematic representation of the variants used as input data in the analysis.** Sankey diagram illustrates the flow and relationships between the variants submitted to ClinVar as of March 2024. The diagram starts with the initial variant summary data accessed through ClinVar FTP site and ends with the variant grouping as Pathogenic, Benign, and Unknown used in the analysis. *focused_Clin_Sig is described as the reported clinical significance of the variants submitted to ClinVar and the specific classes that are focused in the analysis, whereas other_Clin_Sig class represents all the other clinical significance submissions. Sankey diagram was created utilizing a freely accessible tool SankeyMatic, *http://sankeymatic.com/build/.*

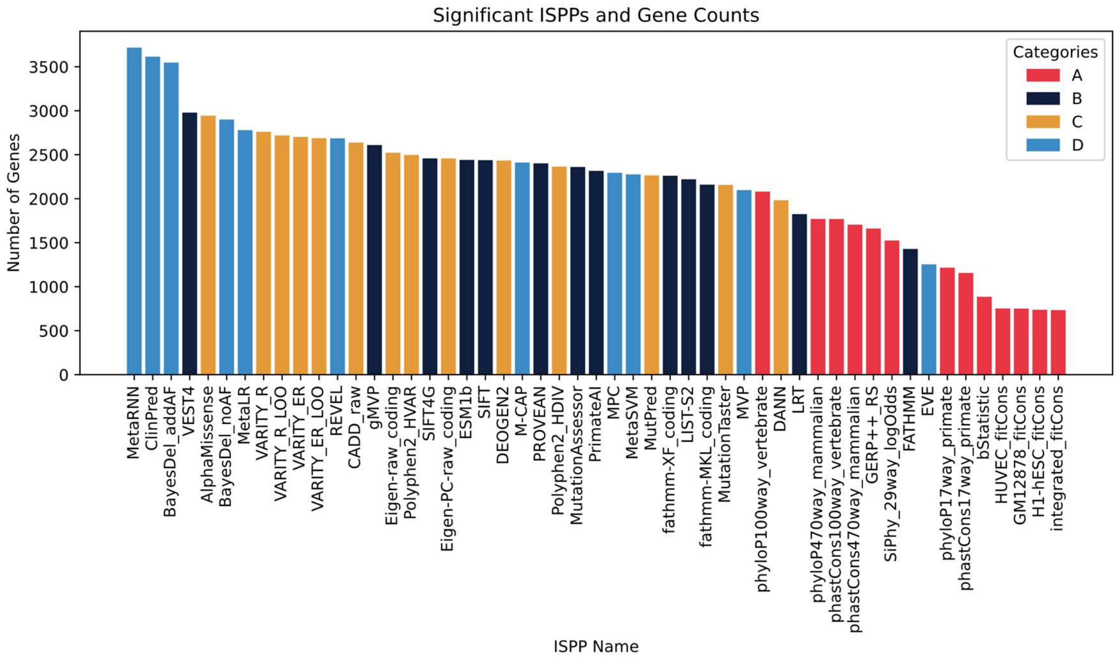

**Supplementary Figure 2. ISPPs and genes with significant results throughout the exome.** The bar plot shows the ISPPs analyzed on the x-axis and gene counts on the y-axis. Gene counts represent the number of genes each ISPP found to be significant in multiple comparisons for every unique gene analyzed in this study, throughout the exome. The bars in the plot are colored based on ISPP categorization, shown in the color legend.

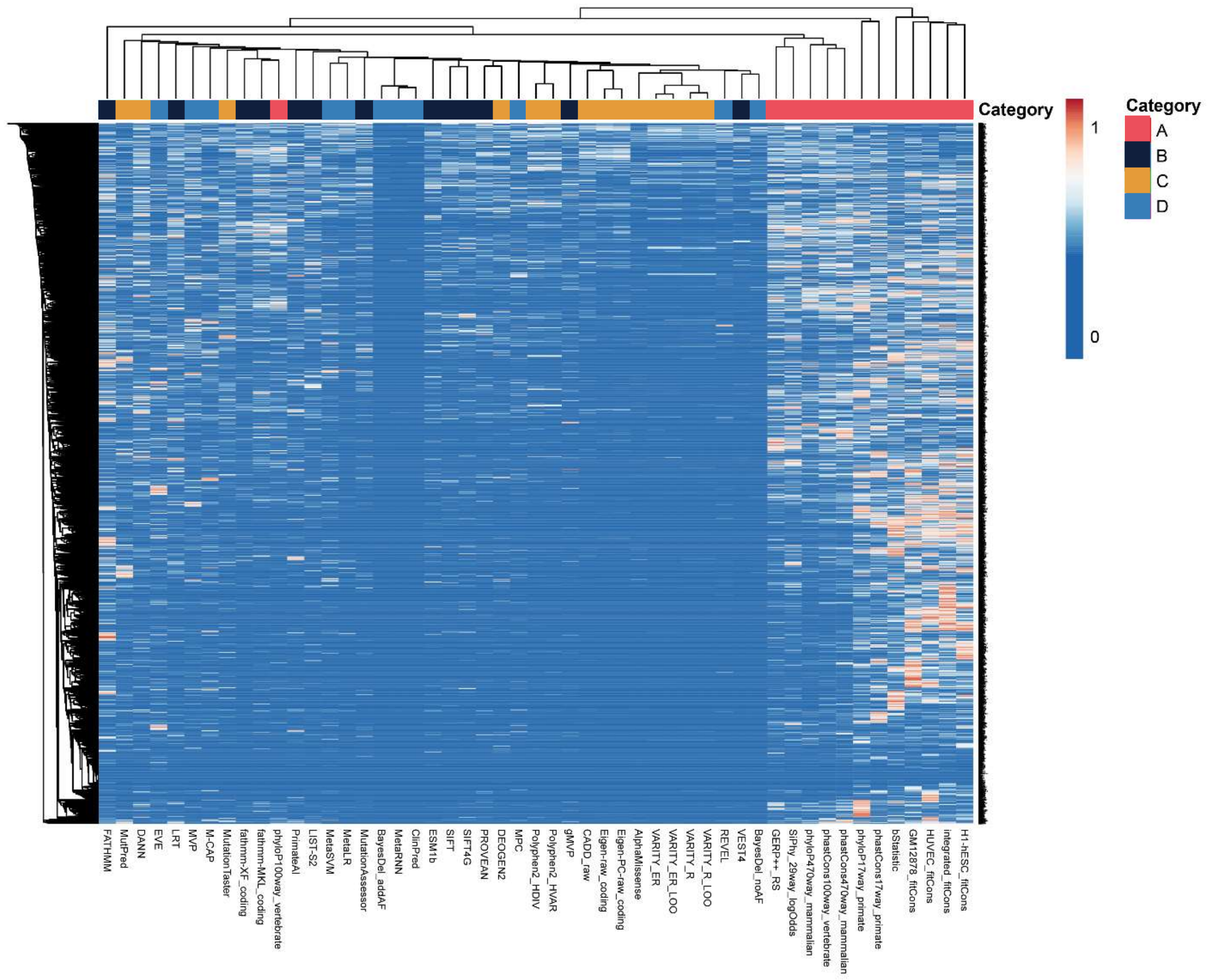

**Supplementary Figure 3. The ISPP correlation heatmap of multiple comparison p-values for the Mendeliome.** The correlation heatmap for Mendeliome of the p-values from multiple comparisons for ISPPs. The dendrogram on the top of the figure shows the hierarchical clustering using the Euclidean distance of the ISPPs, whereas the dendrogram on the left shows the hierarchical clustering result of the Mendeliome genes. ISPP categories are represented by four different colors: red, blue, green, and purple for Category A, B, C, and D, respectively. The color scale for the heatmap was set to be between 0 and 1, ranging from blue to red, which represents the p-values for each gene-ISPP pair. No transformation or scaling was applied to the p-values.

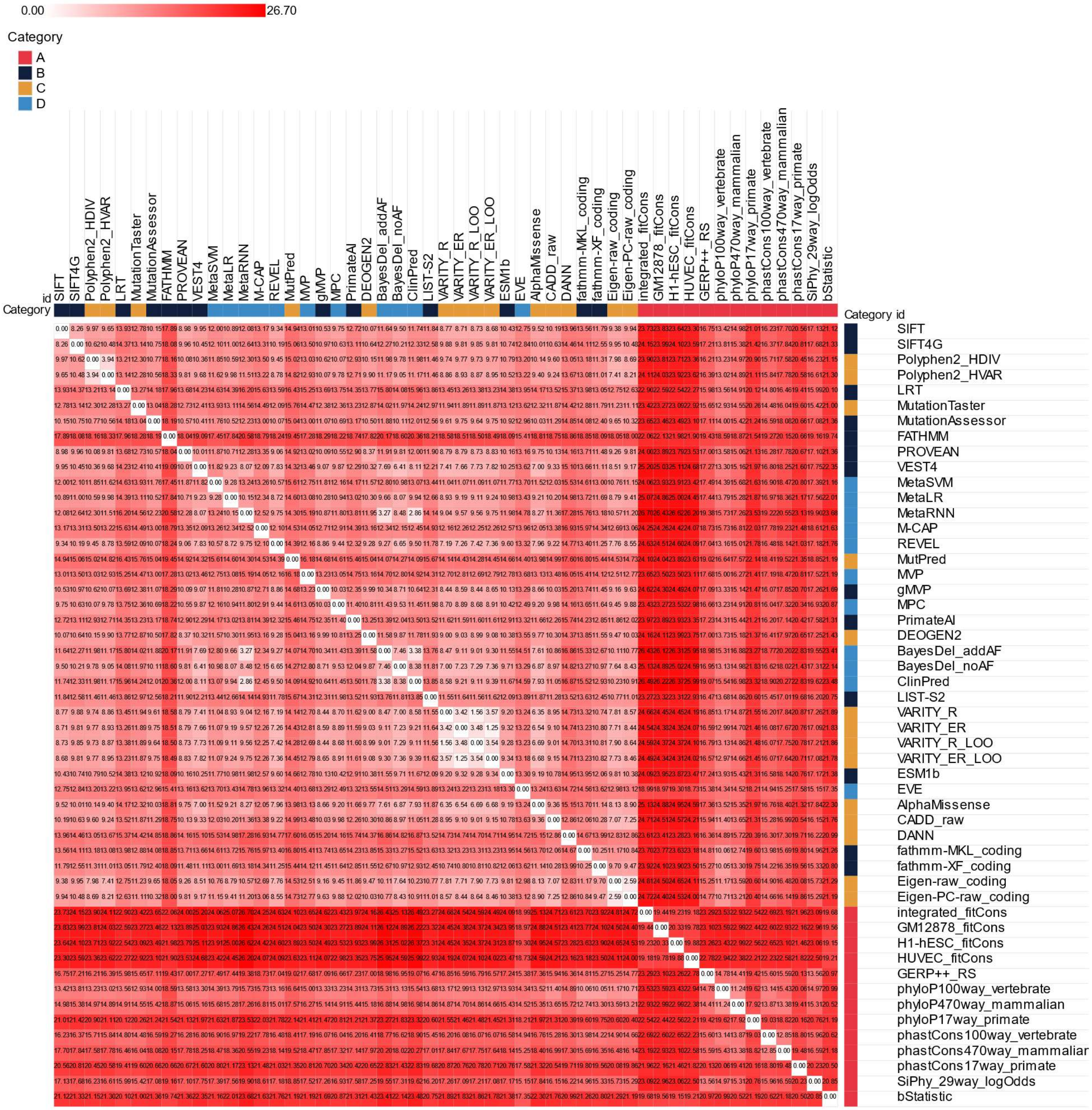

**Supplementary Figure 4. ISPP similarity matrix for the Mendeliome.** The similarity matrix shows the relation between ISPP predictions among Mendeliome genes based on p-values from multiple comparisons utilizing Euclidean distance, shown in each square. The color scale ranges in shades of red, darker shades represent increased distance between two points. ISPP categories are represented by four different colors, shown in legend.

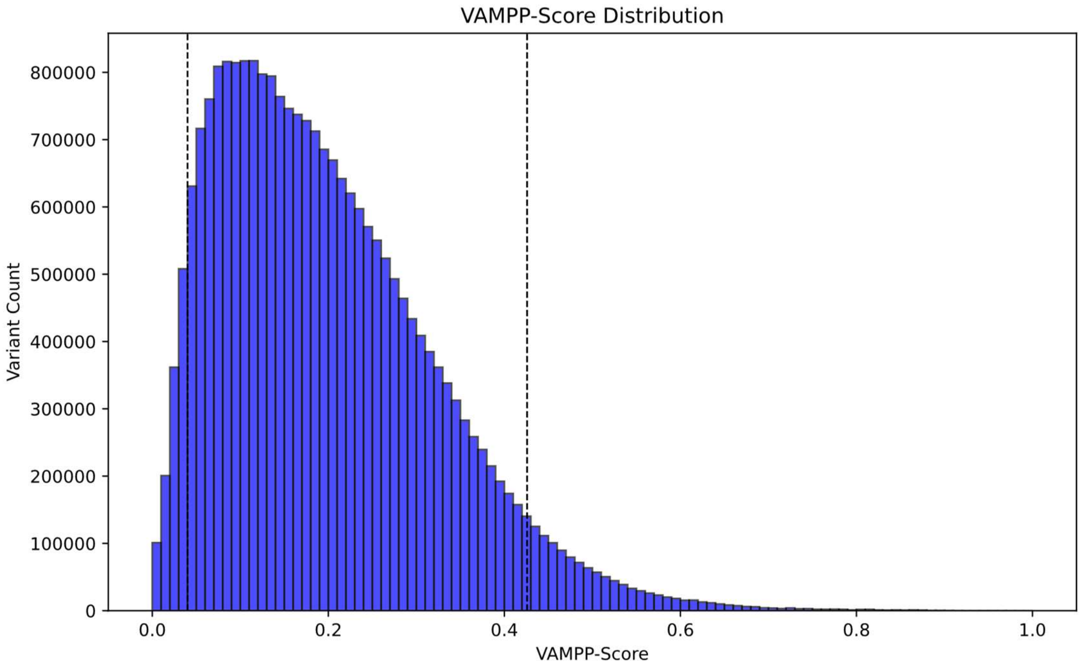

**Supplementary Figure 5. VAMPP-score general distribution.** Histogram shows the distribution of VAMPP-score in a total of 23,464,253 variants from dbNSFP v4.7A (1, 2). The quantiles for 0.05 and 0.95 at score points of 0.04003 and 0.42583, respectively (black dashed line).

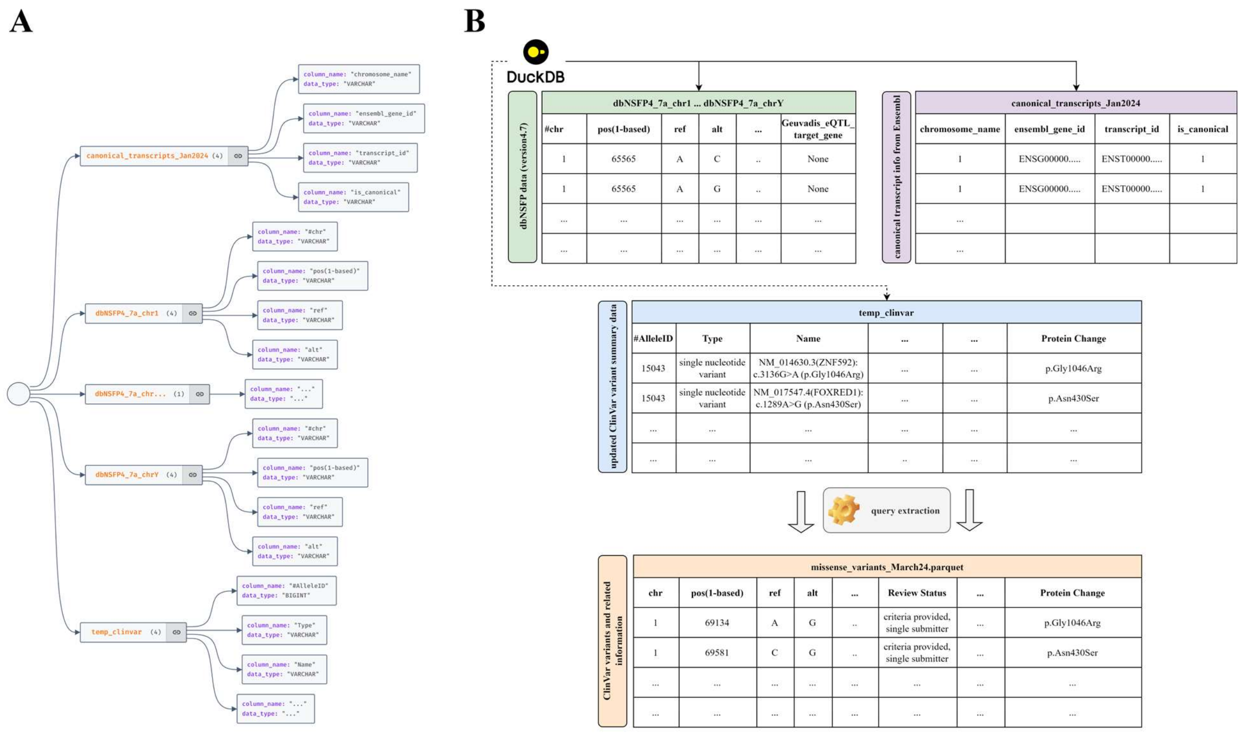

**Supplementary Figure 6.** Schematic representation of the database structure. A. The figure shows the DuckDB (65) database structure. The database mainly consists of twenty-five tables: *canonical_transcripts_Jan2024* and *dbNSFP4_7a* data for each chromosome 1-22, X, and Y. The *canonical_transcripts_Jan2024* table has the canonical transcripts available in the Ensembl.release 111 (29) for the genes in human genome chromosomes, 1-22, X, and Y. *dbNSFP4_7a* chromosome tables include all the data provided in dbNSFP v4.7A (1, 2). *temp_clinvar* represents the ClinVar (3) variant summary data that is updated weekly, which can be reached through the ClinVar FTP site. All the data available in the ClinVar variant summary is integrated as a temporary table, hence named temp_clinvar, which is then used for the variant list querying and matching variant information among the tables. B. The figure shows the general structures of the input and output datasets. The three tables located on the top of the figure, before query extraction, correspond to the database tables shown in A. The extracted file consists of the matching variants from ClinVar with their corresponding ISPP rank scores for the statistical analysis and further downstream applications.
